# Aligning sequence from molecular inversion probes

**DOI:** 10.1101/007260

**Authors:** Brent S. Pedersen, Elissa Murphy, Ivana V. Yang, David A. Schwartz

## Abstract

**Summary:** Molecular inversion probes (MIP’s) allow efficient enrichment of genomic regions of interest for the purpose of targeted sequencing. To date, there is a paucity of simple-to-use software to align sequences derived from this method. Here, we describe a single program that performs mapping, arm removal, and deduplication before outputting alignments in SAM format.

**Availability:** *bwa-mips* is available at https://github.com/brentp/bwa-mips under the MIT license.

## 1 INTRODUCTION

While sequencing costs have dropped dramatically, it is still prohibitive to perform entire genome sequencing on large numbers of samples. Methods such as the use of MIPs for target enrichment can reduce costs and focus efforts on genomic locations likely to yield findings of interest. In addition, the use of MIPs obviates common hurdles of standard library construction resulting in a highly scalable and cost effective method for targeted sequencing across large cohorts. MIP capture is a relatively quick and easy method involving annealing, gap-fill and ligation, followed by exonuclease treatment and PCR amplification all in a single tube. The entire procedure can be completed in two days with minimal hands-on time.

While the sample preparation steps are well-documented, processing the data is non-trivial. Once sequenced and aligned, the synthesized arms must be removed so that synthesis errors are not mistaken for genetic variation in a sample.

## 2 APPROACH

We developed a simple-to-use application that uses BWA mem (Li, 2013) to map the sequence reads, then strips the ligation and extension arms and marks duplicate the reads using a unique molecular identifier (UMI). The application is written in python; it outputs alignments in BAM format (Li *et al.*, 2009) and generates a report that includes the target enrichment and percent of on-target reads.

## 3 METHODS

*bwa-mips* depends on *bwa mem* (Li, 2013) and Picard tools (http://picard.sourceforge.net/). It accepts paired-end fastq files with an optional unique molecular identifier (UMI) at the start of the second read of each pair. Before streaming the sequences to *bwa mem*, *bwa-mips* moves the UMI into the read name so that it can be used later to mark duplicate reads. Once *bwa mem* has aligned the reads, *bwa-mips* removes the ligation and extension arms from the reads guided by the MIP’s design file which contains the columns: ext probe start, ext probe stop, lig probe start and lig probe stop and chr. This matches the output of MIPgen, a recent piece of software to design MIPs (Boyle *et al.*, 2014). The strand and order (first or second) of the read are used to lookup the MIP in the design file. Reads not found to be associated with a MIP are marked with a flag indicating that they failed QC, but they are still reported. Reads associated with a MIP have their sequence, quality, and cigar fields adjusted to remove arms so that synthesis errors are not present in the final aligned sequence. The steps are performed in this order because the read must be mapped before it is possible to know which MIP it belongs to.

The software accounts for insertions and deletions when adjusting both the sequence and the cigar string during arm removal and it adds several tags (Li *et al.*, 2009) indicating the MIP index and the original sequence. Finally, if UMI’s were added during the library preparation, the software marks duplicate reads. Of pairs with the same positions and UMIs, all but the one with the highest mapping quality are marked as duplicates.

Once the alignment is completed, a report indicating the percent of on-target reads and the fold-enrichment (relative to what is expected by chance given the size of the target regions relative to the size of the genome) is shown.

The pipeline is enumerated in Table 1

**Table 1.** steps in bwa-mips.

| step | explanation |
| --- | --- |
| move UMI | UMI is moved from the start of the 2nd read of each pair to the read-name and reads are streamed to *bwa mem* |
| alignment | *bwa mem* is used to align reads |
| MIP lookup | the location and strand of the reads are used to lookup the MIP from the design file |
| arm removal | positions in the design file inform trimming of the alignments such that arms (with potential synthesis errors) are not included |
| tagging and cigar | the cigar string is adjusted to mirror the trimming and SAM tags are added indicating the MIP that each alignment belongs to. |
| mark duplicates | alignments with the same start and UMI are marked as duplicates |
| report | summary report of alignment statistics is generated |

### 3.1 Example Usage

Included in the *bwa-mips* repository is a full example. It includes a MIP design file for a region of chromosome 6 that is compatible with the output from MIPgen (Boyle *et al.*, 2014). Once the reference genome is indexed with *bwa*, the entire pipeline can be run as:

~~~
python bwamips.py ref.fasta mips-design.txt \
        --picard-dir picard-tools-1.117/ \
        $FQ1 $FQ2 > output.sam
~~~

Where *$FQ1* and *$FQ2* are the paired end FASTQ files. The *output.bam* file will contain the alignments in BAM format. For this selected dataset, contained in the *example/* directory of the *bwa-mips* repository, the output report indicates a 3000-fold enrichment of reads to the target region with 86% of reads mapped on-target.

## 4 DISCUSSION

While traditional enrichment procedures are well established, the requirement of mechanical fragmentation and size selection create bottlenecks as sequencing projects are scaled to thousands of samples (Shendure and Ji, 2008). Additionally, due to the upfront library construction, the cost structure of these procedures does not change much whether capturing 100kb or 6Mb. Together these show the importance of software to align MIP data.

While there has been much work to date on designing (Stenberg *et al.*, 2005; Boyle *et al.*, 2014) and using molecular inversion probes (Hardenbol *et al.*, 2003, 2005), there are few tools that will map the resulting data. We have introduced *bwa-mips* to fill this volid.

## 5 CONCLUSION

We have introduced *bwa-mips*, described its implementation and demonstrated its usage on an example dataset.

## ACKNOWLEDGEMENT

Thanks to Evan Boyle and Jay Hesselberth for helpful discussions.

### Funding

Text Text Text Text Text Text Text Text.

